# Magnetic Bead-Based Separation (MBS) of Pneumococcal Serotypes

**DOI:** 10.1101/2021.06.06.447277

**Authors:** Anna York, Emily T. Huynh, Sidiya Mbodj, Devyn Yolda-Carr, Maikel S. Hislop, Haley Echlin, Jason W. Rosch, Daniel M. Weinberger, Anne L. Wyllie

## Abstract

The separation of pneumococcal serotypes from a complex polymicrobial mixture may be required for different applications. For instance, a minority strain could be present at a low frequency in a clinical sample, making it difficult to identify and isolate by traditional culture-based methods. We therefore developed an assay to separate mixed pneumococcal samples using serotype-specific antiserum and a magnetic bead-based separation method. Using qPCR and colony counting methods, we first show that serotypes (12F, 23F, 3, 14, 19A and 15A) present at ∼0.1% of a dual serotype mixture can be enriched to between 10% and 90% of the final sample. We demonstrate two applications for this method: extraction of a known pneumococcal serotype from saliva samples and efficient purification of capsule switch variants from experimental transformation experiments. Moreover, this method may have further laboratory or clinical applications when the selection of specific serotypes is required.

## INTRODUCTION

*Streptococcus pneumoniae* (pneumococcus) is an opportunistic pathogen that resides asymptomatically in the upper respiratory tract of many healthy adults and children worldwide. This asymptomatic colonization is a pre-requisite for the development of pneumococcal disease, including upper respiratory tract infections (such as otitis media), lower respiratory tract infections (such as pneumonia), and invasive pneumococcal disease (IPD) (such as meningitis and bacteremia). Pneumococcal disease often occurs in the very young, elderly, or immunocompromised^1^. Pneumococcus is a leading cause of lower respiratory disease, and contributed to 1,189,937 deaths globally in 2016.^2^

The capsular polysaccharide (CPS) is the outermost layer of encapsulated strains of *S. pneumoniae*, and more than 100 antigenically distinct serotypes have been identified.^3^ Pneumococcal conjugate vaccines (PCV) are highly effective against pneumococcal disease but only cover up to 20 of these serotypes. While pneumococcal disease declined following the introduction of PCVs, a concomitant increase in disease caused by non-vaccine serotypes occurred. This emergence of non-vaccine serotypes in carriage and invasive disease is called serotype replacement.^4^ Serotype replacement occurs for two reasons, first the opening of a new niche in which existing strains expressing capsules not targeted by the vaccine can thrive. Second, vaccine-targeted strains can acquire the capsule biosynthesis cassette from a different serotype, allowing them to evade vaccine-induced immunity. Serotype switching occurs when the *cps* locus from one *S. pneumoniae* serotype (or related species) is transferred into the genetic backbone of another *S. pneumoniae* serotype by transformation.^5^ Genetic exchange between two *S. pneumoniae* serotypes requires co-colonization of two or more serotypes.

In addition to naturally occurring serotype switches,^6–8^ researchers have been generating *cps* switch mutants in the lab for nearly 100 years. The first capsule switch experiments conducted by Griffith in 1928, were accomplished by mixing avirulent, unencapsulated pneumococci with virulent, but killed, encapsulated strains, and injecting this mixture into a mouse. The capsule-switched strains could then be isolated from the mouse.^9^ More recently, generating *cps* switch mutants in the lab has been accomplished using various genetic cassettes.^10,11^ These types of studies have permitted the generation of a number of capsule switch mutants, and this allows for detailed experimental evaluation of the relative importance of capsule and genetic background for different phenotypes.^8,12,13^ Current methods for generating capsule-switched variants require the use of selectable markers, are labor intensive, and not easily scalable. Methods that allow for separation of multiple serotypes could eliminate the need for selection pressure altogether or could be used in combination to conduct higher throughput transformations.

There is also a need to isolate individual pneumococcal strains from clinical samples. Nasopharyngeal swabs have long been considered the gold standard sample type for the detection of carriage of *S. pneumoniae*,^14^ but recent studies have demonstrated utility for saliva to improve the detection of carriage in adults.^15,16^ Whilst testing saliva improves the detection of pneumococci when using molecular methods (such as qPCR), it can be challenging for the isolation of live pneumococcal colonies due to the density and diversity of bacteria present in saliva. A method that enables the separation of pneumococci, in a serotype-specific manner, from other species present in saliva would be useful for clinical and laboratory studies alike.

We developed a magnetic bead-based separation (MBS) method which requires no selection markers and can be used to extract live pneumococci, of a known serotype, from a mixture of pneumococci or from clinical samples containing other bacteria (such as saliva).

## METHODS

Figure 1 summarizes the MBS method; briefly, a mixture of serotypes is incubated with antisera pool(s) unique to the desired serotype, then following wash steps is incubated with secondary antibody conjugated to a magnetic bead. The cells are extracted using the automated Kingfisher Flex Purification System and the eluate plated on blood agar plates. Unless otherwise stated a blood agar plate (BAP) comprises Tryptic Soy Agar (TSA) II supplemented with 5% (v/v) defibrinated sheep blood, and are sometimes referred to as ‘plain plates’. BAPs containing the following concentrations of antibiotics/additives for selection were also used: 0.018 μg/mL, 0.036 μg/mL, 0.18 μg/mL and 0.072 μg/mL penicillin, 10 μg/mL gentamycin, 400 μg/mL kanamycin and 800 μg/mL streptomycin with 10% (w/v) sucrose. Unless otherwise stated all overnight incubations occur at 37°C and 5% CO_2_.

**Figure 1.**
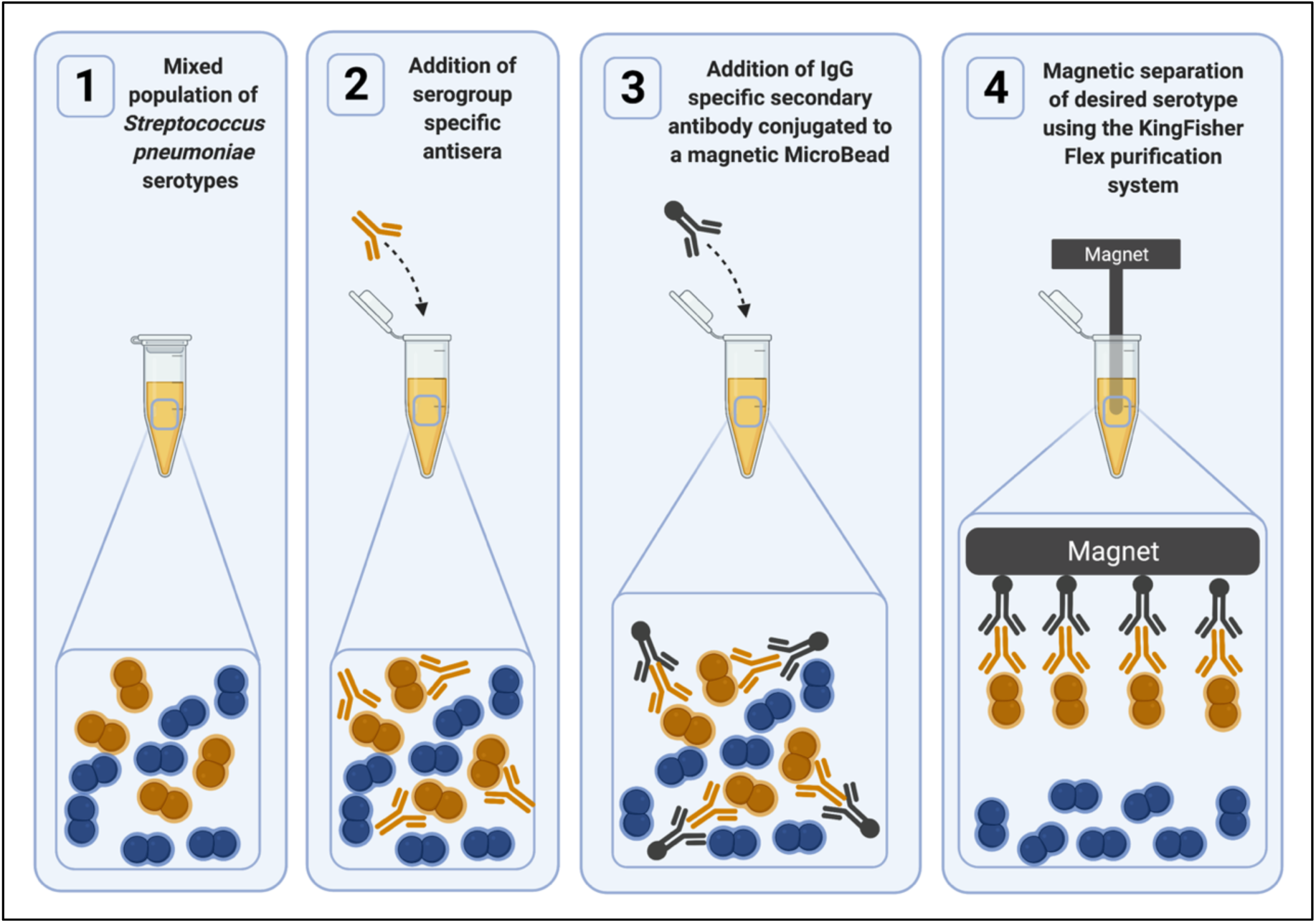
An overview of the Magnetic Bead-based Separation (MBS) method. To a dual serotype mixture (1), antisera specific for the desired serotype are added (2). Following brief wash steps, IgG-specific secondary antibody conjugated to a magnetic bead is incubated (3) and finally the desired cells are extracted using the KingFisher Flex Purification System (4).

### Magnetic Bead-Based Separation (MBS) Method

Approximately 1×10^4^ cells and 1×10^7^ cells from two different serogroups of *S. pneumoniae* were mixed together (∼0.1% minority serotype). Cells were pelleted by centrifugation at 13,000 rpm and resuspended in 450 μl Buffer 1 (1x Phosphate Buffered Saline with 1% bovine serum albumin (BSA)). The resuspended sample was incubated at 4°C on a shaking platform at 150 rpm for 1 hour. The two antisera pools specific for the minority serogroup were combined in a 1:1 ratio and diluted 50-fold in Buffer 1. Next, 30 μl of antisera mix was added to the sample and incubated at 4°C on a shaking platform at 150 rpm for 1 hour. The sample was centrifuged at 13,000 rpm for 5 mins, the supernatant was discarded, and the pellet was resuspended in 450 μl Buffer 1; this step was repeated again. Next, 20 μl of Anti-Rabbit IgG MicroBeads (Miltenyi Biotech) was added, gently vortexed and incubated at 4°C on a shaking platform at 150 rpm for 30 minutes. The sample was extracted using the KingFisher™ Flex Purification System (ThermoFisher) with the protocol detailed in Table S3. The eluted sample was resuspended by pipetting the sample in the elution well 50-100 times before transferring it to a new Eppendorf tube. Following transfer, the sample was thoroughly mixed by vortexing a minimum of 10 times for 5-10s with 5 sec intervals.

To minimize cell losses, when supernatant was removed from cell pellets, 50 μl of supernatant was always left on top of the pellet. The specific rabbit antiserum pools (SSI Diagnostica, Hillerød, Denmark) used for the MBS method, and the SSI ImmuLex™ Pneumotest Pools used for serotyping are outlined in Table S1.

### Proof of concept and primary analysis

To demonstrate proof of concept for the MBS method we used three pairs of six different serotypes where one serotype in each pair was penicillin resistant and the other penicillin sensitive. It is important to note that different penicillin sensitivity is not necessary for separation but was instead used to make the quantification of the efficiency of this method easier. The three pairs were 12F and 23F (Pair 1), 3 and 14 (Pair 2) and 19A and 15A (Pair 3). Serotype 3 exists as two distinct morphologies; small non-mucoid colony variant (SCV) and mucoid variant.^17^ We therefore isolated SCV and mucoid variants and chose to work primarily with the SCV for three reasons; SCVs are easier to count, easier to isolate as single colonies (for serotyping) and less easy to distinguish from other serotypes based on morphology, thus reducing selection bias during the colony selection for serotyping. The MIC of each serotype was determined using penicillin E-strips, and then the exact concentration of penicillin for blood agar plates was determined experimentally by varying the penicillin concentration and plating out cells at known CFU/mL. The concentration of penicillin used in the blood agar plates was the concentration at which the resistant serotype grew equally well on a penicillin containing plate, as it did on a plain plate, whilst the susceptible serotype showed no growth on the penicillin containing plate but normal growth on a plain plate. For Pairs 1, 2 and 3, BAPs containing 0.018 μg/mL, 0.036 μg/mL and 0.18 μg/mL penicillin were used, respectively.

For all three pairs, Sample R is when the penicillin resistant serotype is the minority species, and Sample S is when the penicillin sensitive serotype is the minority species. Samples were plated out onto BAPs with and without penicillin, at two stages in the protocol; immediately prior to the first incubation (PRE), and after extraction (POST). In all cases 5 μl of sample was serially diluted in 45 μl PBS, in triplicate. For samples where the minority strain was penicillin resistant, 20 μl of sample at a 10^−1^ dilution was plated on penicillin plates, while 20 μl of sample at a 10^−4^ dilution was plated on plain blood agar plates. In samples where the majority serotype was penicillin resistant, 20 μl of sample at a 10^−4^ dilution was plated on both BAPs with and without penicillin. In addition to the diluted samples, 10 μl of undiluted sample at the PRE and POST stage, and the remaining volume (∼40 μl) after elution was plated on BAPs, to provide DNA for qPCR experiments conducted to establish separation efficiency. In all cases 10 μl or 20 μl samples were pipetted onto the BAP and the plate was then tilted to allow the sample to run down the length of the plate. The BAPs were incubated overnight.

### Secondary analyses

To establish if separation efficiency was similar for both mucoid (Muc) and single colony variants (SCV) of Serotype 3, two additional pairs; 23F and 3SCV (Pair 4), and 23F and 3Muc (Pair 5) were investigated. These experiments were conducted in duplicate, and efficiency assessed by colony counting and qPCR methods. Pair 4 and 5 used BAPs containing 0.072 μg/mL penicillin.

To investigate the effect of initial proportion of minority serotype on the efficiency of separation, 23F and 12F (Pair 1) were again used. The initial amount of majority serotype (12F) was kept constant at 1×10^7^ CFUs, while the minority serotype (23F) was varied (5×10^4^, 1×10^4^, 5×10^3^ and 1×10^3^). These experiments were conducted once for each dilution, and efficiency was assessed by colony counting and qPCR methods.

The experiments above were conducted using two pooled antisera that were specific for the minority serotype. We investigated whether a single pool of antisera could also be used. This is important because certain pairs of serotypes can only be distinguished by one pool. Serotype pairs which could not be distinguished based on penicillin sensitivity (and therefore could not be assessed by colony counting methods), were used for this analysis, and for pairs which shared a common antisera pool, only the unique antisera was used. These experiments were conducted once for each condition, and efficiency was assessed by qPCR alone.

### Colony counting to quantify separation efficiency

Colonies were counted and the mean colony number was determined, which was then used for downstream analysis. The following equations for Sample R and Sample S were used to determine the percentage of the minority serotype present at each time point.

### Sample R Equation

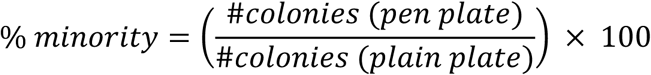

### Sample S Equation

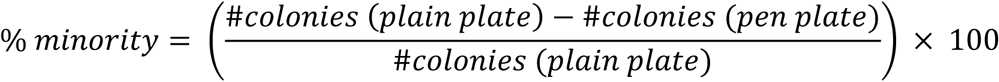

### Serotyping of colonies to confirm separation efficiency

Eight colonies were picked at random from the plain blood agar elution plates and expanded to create a lawn on 1/8^th^ of a BAP and incubated overnight. The serotype of each lawn was confirmed by testing each of the four antisera pools specific to both the majority and the minority serotype in the pair, using ImmuLex™ Pneumotest (SSI Diagnostica) reagents.

### Real-Time qPCR to confirm separation efficiency

Colonies/lawns from each sample, grown on BAP, were harvested into 200 μl PBS using a cotton swab and the DNA was extracted using a DNeasy Blood and Tissue Kit (QIAGEN) as per the manufacturers protocol. DNA concentration was measured using Qubit™ as per the manufacturers protocol. A no-template negative control was included for each primer pair used, ^18^ and a standard curve (positive control) was constructed using genomic DNA from each of the six serotypes under investigation. Each qPCR reaction was 25 μl total volume, consisting of iQ™ SYBR® Green Supermix (BioRad), 5 μl of template DNA and 200 nM of each primer. The real time qPCR was run on a BioRad CFX96™ Touch Real-Time qPCR System. The cycling conditions were 1 cycle of denaturation at 95°C for 10 minutes, followed by 40 cycles of 95°C for 15 seconds and 60°C for 1 minute for amplification, and a melt curve from 65°C to 95°C in increments of 0.5°C. For each sample, amplification with primer pairs from both the minority and majority serotype was conducted in duplicate, the mean of duplicates was used for downstream analysis. The concentration of each serotype in a sample was determined by comparing the C_T_ value to the standard curve for the corresponding serotype.

### Applications for MBS method - Generating capsule-switch mutants by transformation

To determine whether the MBS method could be used to improve capsule switching experiments (by reducing workload and scaling-up transformations), genomic DNA (gDNA) from four donor serotypes (12F, 23F, 35B, 11B) was transformed individually and as a mixed sample into the recipient D39SΔcps::SweetJanus.^11^ The mixed sample was processed with and without the use of the MBS method. An individual transformation of D39 gDNA into D39SΔcps::SweetJanus was included as a positive control.

With the exception of using Todd Hewitt supplemented with 0.5% Yeast Extract (THY) media for liquid cultures, gDNA was extracted as outlined previously.^11^

Frozen stocks of D39SΔcps::SweetJanus were inoculated onto BAP and incubated overnight. Cells harvested from the BAP were used to inoculate THY media to a starting OD_620_ of 0.04 AU, and were grown at 37°C and 5% CO_2_ until late logarithmic phase (OD_620_ ∼0.80). For each of the five individual transformations, 1 mL of culture was transferred into a 1.5mL Eppendorf tube, 3 µg/mL of competence stimulating peptide 1 (CSP1) and 4 μg of the appropriate DNA was added. For the mixed transformation, 4 mL of culture was transferred to a 15 mL falcon tube, 3 µg/mL CSP1 and 4 µg of each of the four gDNA templates was added. Cells were incubated for three hours at 37°C. Subsequently, individual transformation and mixed transformation samples were positively selected for by plating on BAP supplemented with 800 μg/mL streptomycin and 10% (w/v) sucrose (Strep/Suc plates), and incubated overnight.

For the five samples that underwent individual transformations, eight colonies each were selected and expanded onto new Strep/Suc plates and incubated overnight. These expanded samples were re-plated onto both Strep/Suc plates, as well as BAP supplemented with 400 μg/mL kanamycin (Kan plates), for negative selection, and incubated overnight. Colonies that grew on Strep/Suc but not Kan plates were serotyped to confirm they have successfully gained the capsule.

For the mixed transformation sample, all colonies were harvested using a cotton swab and resuspended in 1.5 mL Brain Heart Infusion (BHI) media + 10% (v/v) glycerol. As a control, 100 μl of the mixed sample was serially diluted to 10^−6^, then 100 μl of 10^−4^, 10^−5^ and 10^−6^ dilutions were plated on BAP, and incubated overnight. Following, 100 μl of the mixed sample was aliquoted into four 1.5 mL Eppendorf tubes, centrifuged at 13,000 rpm resuspended in 500 μl Buffer 1 and processed through MBS using the appropriate antisera pool(s) for targeting the appropriate serotype. The elution was plated on BAP and incubated overnight. Thirty-two colonies were selected from the mixed sample that did not undergo MBS, and eight colonies were selected from each of the four samples that had undergone MBS. The serotype of all expanded colonies was determined using SSI latex agglutination.

### Applications for MBS method - Isolating pneumococci from saliva

De-identified pneumococcus-negative saliva samples were obtained from healthy volunteers (< 30 years of age; IRB protocol number 2000029374). De-identified pneumococcus-positive saliva samples were obtained from healthy volunteers (> 60 years of age; IRB protocol number 2000026100). The relationship between qPCR cycle threshold (C_T_) value and CFU/mL was determined using pneumococcus-negative saliva, spiked with pneumococci (serotype 19A) at a variety of known CFU/mL. The concentration of the 19A stock was determined to be 5×10^9^ CFU/mL, which was then serially diluted 1:10 in pneumococcus-negative saliva. After two hours at room temperature, 100 μl of each sample was plated onto BAP supplemented with 10 μg/mL gentamycin (Gent plates) and incubated overnight. The lawn of each culture-enriched saliva sample was harvested into 2100 μl BHI + 10% (v/v) glycerol using an L-shaped spreader. gDNA for each sample was extracted using a standard protocol (Table S4), all DNA templates were tested by qPCR for the pneumococcal gene *piaB* using Luna® Universal One-Step RT-qPCR mix, 2.5 µl template DNA and 200 nM of each primer and probe (Table S2) in a total reaction volume of 20 μl. The cycling conditions were 1 cycle of denaturation at 95°C for 3 minutes, followed by 40 cycles of 98°C for 15 seconds and 60°C for 30 seconds. C_T_ values were plotted against CFU/mL of 19A in the raw saliva sample (Figure S1).

Using data from Figure S1 in combination with data from previous studies^19^ we were able to determine suitable concentrations for spiked-saliva, that reflect levels commonly found in saliva obtained from the healthy individuals during carriage studies. Pneumococcus-negative saliva was spiked with pneumococci (serotype 19A) at varying concentrations (5×10^4^, 5×10^3^, 5×10^2^ and 5×10^1^ CFU/mL) and left at room temperature for 2 hours. Following, 100 μl of each sample was plated onto Gent plates and incubated overnight. The lawn of the culture-enriched saliva was harvested into 2100 μl BHI + 10% (v/v) glycerol.

From each culture-enriched saliva sample, 10 μl was added to 490 μl Buffer 1, and cell separated using the MBS protocol, with the following modifications. The primary incubation step was conducted using SSI antisera (∼16.8 μg total protein) and SunFire Bio monoclonal antibody (mAb) (∼16.8 μg total protein) combined. The secondary incubation was conducted using ∼48 μg total protein of anti-mouse IgG MicroBeads (Miltenyi Biotech) to target the mAb only. As a negative control, culture-enriched saliva samples did not undergo MBS and were instead serially diluted to 10^−6^, the 10^−4^, 10^−5^ and 10^−6^ dilutions were plated on BAPs and incubated overnight.

Colonies that looked like pneumococci (small, grey, moist colonies with a green zone of alpha-hemolysis), were isolated and expanded onto new BAP: sixteen colonies from each sample (with and without cell separation) that contained 5×10^4^, 5×10^3^, 5×10^2^ CFU/mL of 19A in raw saliva samples, and 24 colonies from the sample (with and without cell separation) that contained 5×10^1^ CFU/mL of 19A in raw saliva. Each expanded colony was optochin tested to confirm whether it was pneumococcus (optochin sensitive) or another oral bacteria (optochin resistant). Where a ring of optochin sensitivity was observed but a second (contaminating) bacteria with optochin resistance was also present or, where satellite colonies of pneumococcus were present within the zone of inhibition, samples were considered ‘pneumococcal colonies’ since pure pneumococci can be isolated from the contaminant.

## RESULTS

The MBS proof of concept experiments showed that for all six serotypes, the minority serotype was successfully enriched from ∼0.1% starting percentage to between 13% (serotype 14) and 90% (serotype 3) post MBS, corresponding to a 100-to-900-fold enrichment (Figure 2a). The final percentage of the minority varied between serotypes but was relatively consistent between the three replicates. There was generally good concordance in the estimated MBS efficiency as determined by the qPCR and colony counting (Figure 2b), however efficiency determined by colony counting seemed to be higher and lower than with qPCR for serotype 14 and 3, respectively. Eight colonies from each elution plate were selected at random and in every single case, minority serotype colonies were identified by serotyping (Table 1). This demonstrates that this technique can be used to recover a desired serotype from a dual mixture.

**Figure 2.**
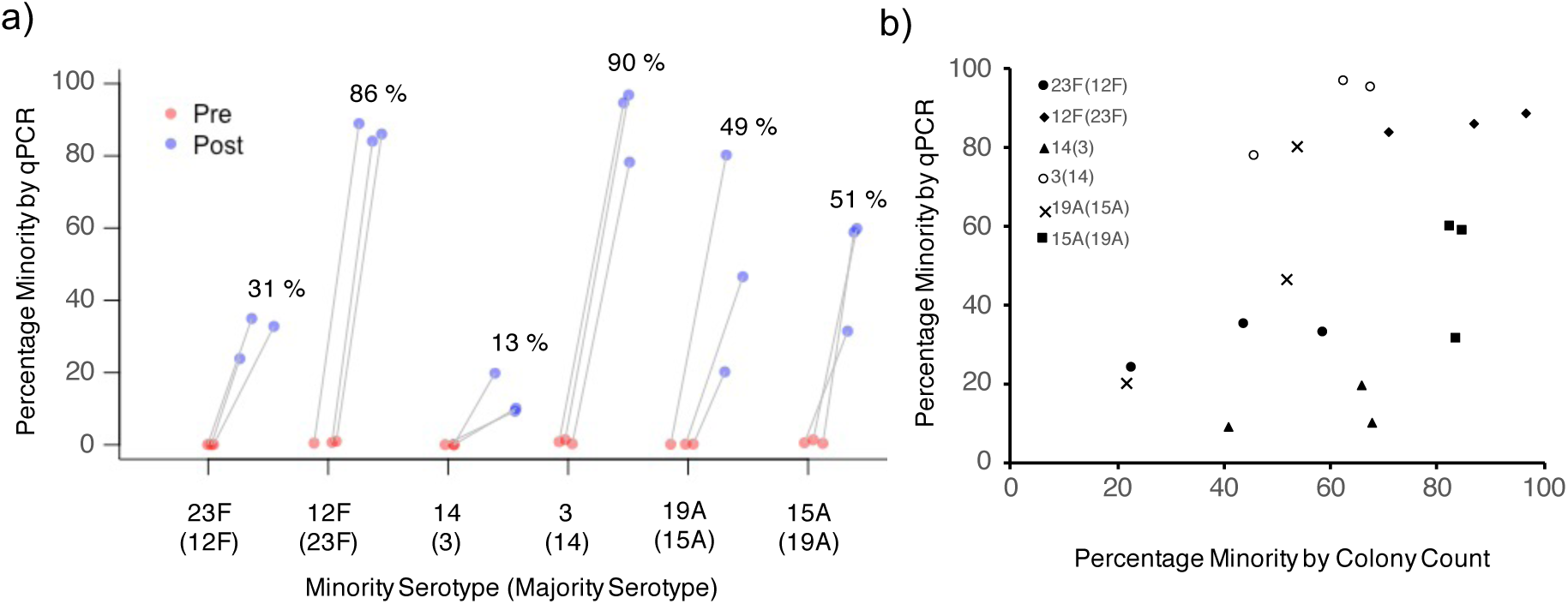
Efficiency of minority strain enrichment using the MBS method on six serotypes. A) Percentage minority serotype present prior to MBS (Pre) and after MBS (Post). The average percentage minority in the post sample is presented above the data points, b) Percentage minority serotype present after MBS (Post) as determined by qPCR and colony counting. Minority and majority serotypes are displayed in the legend in the following format, minority(majority). Results are shown in triplicate for each of three serotype pairs where each serotype of the pair was tested as the minority serotype.

**Table 1.** Total number of colonies on the plain blood agar elution plate (out of eight selected at random) that were positive for the minority serotype by SSI latex agglutination. Positive results reported were those that tested positive with minority serotype antisera and negative with majority serotype antisera.

| Serotype Pairs |  | Number of colonies of the minority serotype following MBS<br>(from eight selected at random from plain blood agar plates) |  |  |
| --- | --- | --- | --- | --- |
| Minority Serotype | Majority Serotype | Rep1 | Rep2 | Rep3 |
| 23F | 12F | 5 | 5 | 5 |
| 12F | 23F | 8 | 6 | 5 |
| 14 | 3 (SCV) | 4 | 4 | 3 |
| 3 (SCV) | 14 | 7 | 6 | 6 |
| 19A | 15A | 1 | 7 | 3 |
| 15A | 19A | 7 | 6 | 5 |
| 23F | 3(SCV) | 4 | 2 | N/A |
| 3(SCV) | 23F | 6 | 8 | N/A |
| 23F | 3(Muc) | 6 | 8 | N/A |
| 3(Muc) | 23F | 5 | 7 | N/A |

A secondary analysis was conducted to identify whether serotype 3Muc was also enriched with a similar efficiency as serotype 3SCV, and to gain insight into how separation efficiency varies when the majority serotype of the pair is altered. MBS was conducted on Pair 4 (23F and 3SCV and Pair 5 (23F and 3Muc). The results were compared to MBS results obtained previously for enrichment of minority serotypes 23F or 3SCV when paired with another majority serotype (namely serotype 12F and serotype 14 from Pair 1 and Pair 2, respectively). The percentage enrichment for both 23F and 3 remained similar even when the majority serotype of the pair was altered (Figure 3). Furthermore, it demonstrates that the MBS method permits successful enrichment of both SCV and mucoid variants of serotype 3, and that the efficiency is similar regardless of the morphology. In all cases minority serotype single colonies were isolated from the elution plate by selection of single colonies and confirmed to be the desired serotype using SSI latex agglutination (Table 1).

**Figure 3.**
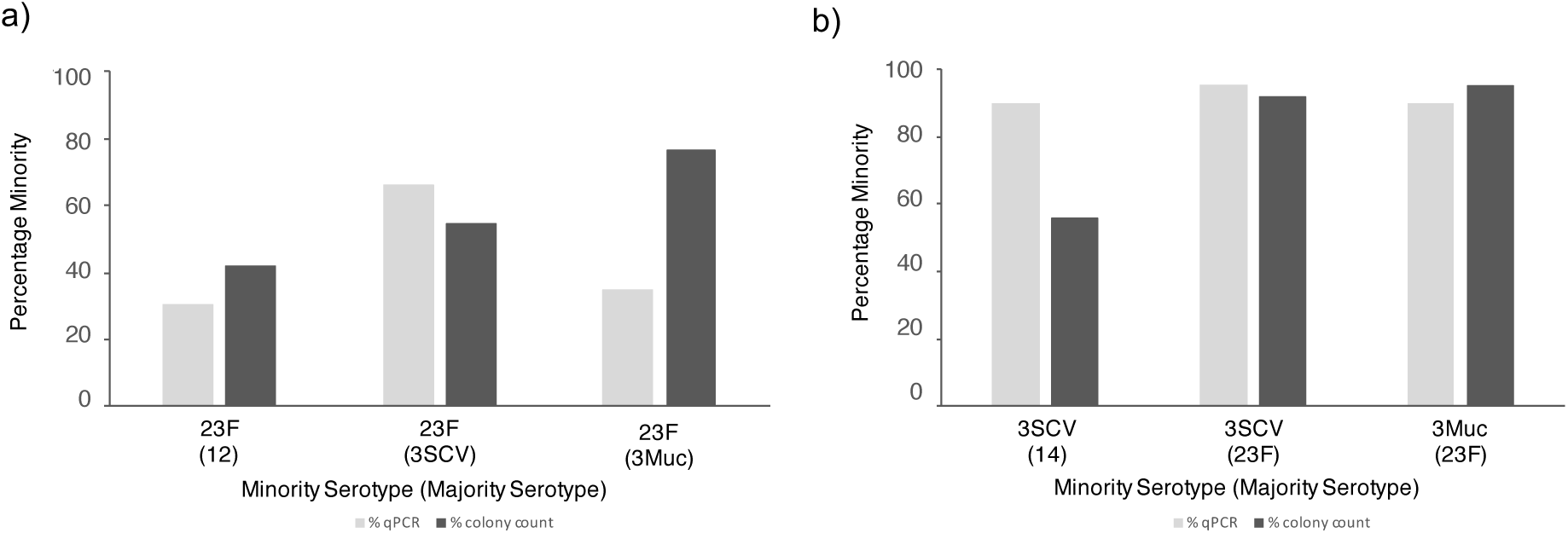
Final percentage of minority serotype after MBS. Averages of triplicate results from Figure 1 are shown for 23F (when with a majority of 12F) and 3 (when with a majority of 14). Averages of duplicate results were plotted for 23F (when with a majority of 3SCV), 23F (when with a majority of 3Muc), 3SCV (when with a majority of 23F) and 3Muc (when with a majority of 23F). Percentage minority from both colony counting and qPCR methods is shown.

The primary analysis specifically used serotype pairs that could be distinguished using two unique pools of antisera. MBS was then tested on eight serotype pairs using only a single antisera pool. A total of six antisera pools (H, P, B, E, R, H and Q) were tested and all were able to successfully enrich a ∼0.1% minority serotype to between 10% and 99% in the final sample (Figure 4a).

**Figure 4.**
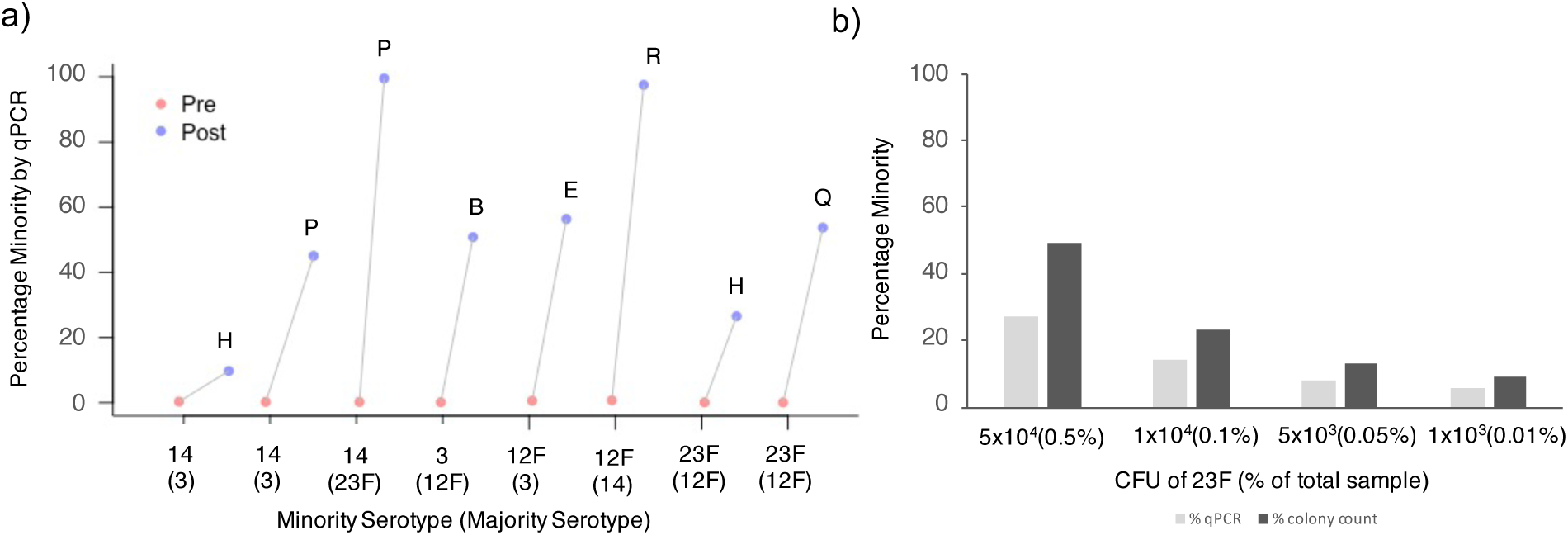
Efficiency of minority enrichment using the MBS method on eight serotypes. A) Percentage minority serotype present prior to MBS (Pre) and after MBS (Post), b) Percentage minority 23F present after MBS (Post) as determined by qPCR and colony counting. Minority and majority serotypes are displayed in the following format, minority(majority). Results shown are singlicate data points only.

Additional analysis aimed to determine whether enrichment was constant at different % minorities. The 23F and 12F pair were used with the majority serotype (12F) remaining constant at 1×10^7^ CFUs and the minority serotype (23F) at four different concentrations in the initial sample. Enrichment of the minority serotype can be achieved even when the starting percentage of a minority serotype is as low as 1×10^3^ CFUs. However, as the initial % minority decreases the percentage minority recovered following MBS also decreases. For initial samples containing 5×10^4^, 1×10^4^, 5×10^3^ and 1×10^3^ CFU’s of minority serotype 23F, the corresponding percentage of 23F present in the final samples were 27%, 14%, 8% and 6% respectively as determined by qPCR, or 49%, 23%, 13% and 9% respectively as determined by colony counting (Figure 4b).

In order to separate serogroups that share reactivity to one antiserum pool, the MBS method should be used with only a single antiserum pool. We therefore investigated outcomes when using one or two antisera Pools and compared the efficiency of antisera pools in the presence of different majority serotypes. MBS of serotype 14 from a majority serotype 3, using both antisera Pool H and Pool P, resulted in the final sample containing ∼13% of serotype 14. However, use of only Pool H or Pool P, at an equal final volume to the combined pools, resulted in serotype 14 being 10% and 45% of the final samples respectively. Therefore, in this example, Pool P alone achieves the greatest efficiency of MBS, but in the absence of knowing which antisera is more efficient, and if the serotype pairs permit dual use, it would be prudent to combine both antisera pools. Furthermore, we confirm that the overall efficiency of enrichment achieved by any antisera pool, is not only dependent upon the minority serotype alone, but also the majority serotype. The final percentage of serotype 14 following MBS (using Pool P) from a majority serotype 23F, is 99%, more than double the percentage of serotype 14 present following MBS (using Pool P) from a majority serotype 3.

### Generating capsule-switch mutants by transformation

For transformations conducted individually, the positive control (D39 gDNA) was successfully transformed into D39SΔcps::SweetJanus at the cps locus, with 8/8 colonies serotyping as serotype 2. gDNA from serotype 23F and serotype 35B was also found to transform into D39SΔcps::SweetJanus at the cps locus, with 7/8 and 8/8 colonies serotyping as 23F and 35B respectively. Conversely, 0/8 colonies were serotyped as 12F or 11B for these transformations, suggesting that transformation may not have occurred or may have occurred at very low efficiency (Table 2).

**Table 2.** Number of positive transformations (per 8 colonies) when individual transformations are conducted using the standard transformation procedure and selection methods.

| Serotype | Individual Transformations<br># of transformants |
| --- | --- |
| D39 | 8/8 |
| 12F | 0/8 |
| 23F | 7/8 |
| 35B | 8/8 |
| 11B | 0/8 |

Transformations conducted in higher throughput (i.e. gDNA from 4 serotypes combined) show that in the absence of cell separation, even when gDNA samples are mixed, it is possible to isolate transformation to 23F and 35B with 9/32 and 11/32 colonized serotyping as these serotypes, respectively. Similarly, to the results seen in the individual transformations, transformants of 12F or 11B were not identified (0/32). The mixed transformations that were subsequently cell separated with MBS to enrich for the desired serotype showed that 23F, 35B and 11B were successfully transformed, with 8/8, 5/8 and 7/8 colonies identified to be 23F, 35B and 11B, respectively (Table 3). This confirms that 11B is able to transform into D39SΔcps::SweetJanus at the cps locus but this likely occurs at a lower efficiency, or that acquisition of the *cps* locus requires additional non-cps recombination from other serotypes in the mix. For serotype 12, colonies were observed on the BAP following MBS, however 0/8 were identified to be 12F transformants, therefore this transformation may only occur at very low frequencies, under very specific conditions, or not at all. Of the 8 colonies selected from the cell separation enriching for 12F, 6/8 were serotype 23F, 1/8 were serotype 11B and only 1/8 was untransformed. For serotype 35B, 1/3 was serotype 23F and 2/3 were untransformed. For serotype 11B, 1/8 was serotype 23F. The presence of these contaminating serotypes suggests that the antisera/antibodies used in MBS have some non-specific cross-reactivity.

**Table 3.** Number of positive transformations for each serotype from a mixed transformation containing gDNA of 12F, 23F, 35B and 11B, in the presence of cell separation (8 colonies picked per sample) and in the absence of cell separation picked (total of 32 colonies picked).

| Serotype | Control (no cell separation)<br># of transformants | Cell Separation<br># of transformants |
| --- | --- | --- |
| 12F | 0/32 | 0/8 |
| 23F | 9/32 | 8/8 |
| 35B | 11/32 | 5/8 |
| 11B | 0/32 | 7/8 |

### Isolating pneumococci from saliva

To determine if the MBS method could be used to enrich for a known serotype in pneumococcus-positive saliva, we spiked two saliva samples (A and B) which tested qPCR-negative for pneumococcal genes *piaB* and *lytA*, with varying concentrations of serotype 19A and compared the success of identifying pneumococcal colonies in the presence and absence of MBS (Table 4). For both saliva A and saliva B, at all concentrations of 19A, the MBS method resulted in equal or improved isolation of pneumococcal colonies. In saliva A, the MBS method was still able to enrich for pneumococcus when the concentration of 19A was 5×10^1^ CFU/mL in raw saliva, however for Saliva B the MBS method was only successful at a 19A concentration of 5×10^3^ CFU/mL in raw saliva. The sensitivity of this assay is therefore dependent upon not only the concentration of pneumococci in the sample but also the composition of saliva itself, and may vary from sample to sample. The MBS method was then tested on a saliva sample that had tested qPCR-positive for serotype 15B/C but from which we had been unable to isolate pneumococcus using the standard culture-based dilution method.^20^ Here, the MBS method successfully enriched for pneumococcus in the sample, and of the 32 colonies selected, 29 were confirmed to be pneumococcus.

**Table 4.** Isolation of optochin-sensitive pneumococcal colonies using the standard dilution method compared to the MBS method. Two saliva samples (A and B) were spiked with four concentrations of serotype 19A. Pure pneumococcal colonies were identified by a zone of inhibition around the optochin disk, any colonies that were mixed colonies (i.e. those with a zone of inhibition but some secondary growth (a non-pneumococcal contaminant) growing within the zone of inhibition, or, had satellite colonies appearing withing the zone of inhibition) were considered to be successful isolation of pneumococcus).

| CFU/mL of *S. pneumoniae* 19A | # of 19A colonies identified in Saliva A |  | # of 19A colonies identified in Saliva B |  |
| --- | --- | --- | --- | --- |
|  | With MBS method | Standard dilution method | With MBS method | Standard dilution method |
| $5 \times 10^4$ | 15/16 | 15/16 | 15/16 | 1/16 |
| $5 \times 10^3$ | 15/16 | 5/16 | 5/16 | 0/16 |
| $5 \times 10^2$ | 16/16 | 2/16 | 0/16 | 0/16 |
| $5 \times 10^1$ | 6/24 | 0/24 | 0/24 | 0/24 |

## DISCUSSION

We developed the MBS method that can enrich for a desired serotype from a mixed-serotype sample in a laboratory setting. Enrichment using the MBS method was demonstrated for six serotypes (23F, 12F, 3, 14, 15A and 19A) including two serotypes with more unique capsules (serotype 3 and serotype 14). We were able to demonstrate two use cases for this method: separation of capsule switch mutants (from mixed transformation experiments) and, enrichment of pneumococcus from saliva samples.

In the primary analysis used to develop the MBS method, we show that all six of the minority serotypes investigated (23F, 12F, 14, 3, 19A and 15A) can be successfully enriched from ∼0.1% of an initial mixed serotype sample to up between 13% and 90% in the final sample. The inclusion of Serotype 3 (which exists as single colony and mucoid variants) and serotype 14 (which has an uncharged capsule) in this panel, showed that this method is suitable for serotypes with more unique capsules. Two methods; colony counting, and qPCR were employed in order to assess efficiency of the MBS method. The estimates from both methods were broadly concurrent but there are a few examples where the efficiency estimates do differ. This may be explained by the formation of varying chain lengths in pneumococcus, such that if the two serotypes in a pair form vastly different length chains, the estimations of efficiency may be biased. A serotype that readily forms chains would result in an underestimation of its presence in the sample using the colony counting method, but qPCR would provide a more accurate estimation. Despite some differences in efficiency estimates between colony counting and qPCR methods, we were able to successfully isolate minority serotype colonies post MBS in all cases. This demonstrates a tangible utility for this method in the laboratory setting. When separating a mixture of cells only a small number of colonies must be isolated to identify the desired serotype. This method therefore allows for the easy recovery of serotype-specific *S. pnuemoniae* isolates.

In the secondary supporting analysis, we compared how enrichment of a minority serotype varied when in the presence of different majority serotypes. A minority serotype 23F was paired with a majority serotype of either 12F or 3, and minority serotype 3 was paired with a majority serotype of either 14 or 23F. With minority 23F, some variation in efficiency of MBS was noted when the majority serotype was changed, however for minority serotype 3, the enrichment efficiency remained very similar despite the change in majority serotype pair. This suggests that the serotype with which the minority is mixed may have some impact on the efficiency of MBS, but it is likely primarily determined by the avidity of the antisera for the desired serotype. Unlike the majority of pneumococcal serotypes, serotype 3 utilizes the synthase-dependent pathway for CPS production, resulting in non-covalently bound CPS which can be released from the glycolipids or synthase.^21^ The CPS of serotype 3 is not covalently linked to the peptidoglycan and can be released,^22^ which leads to a reduction in the protective effect of anti-Type 3 CPS antibodies induced by the PCV13,^23^ we were therefore surprised to find that the MBS method can successfully extract serotype 3 from a mixed sample. This success may be explained by the fact that the cells are not actively growing and likely therefore not releasing CPS into the environment. Furthermore, it is intriguing but reassuring that the efficiency of enrichment between mucoid and SCV serotype 3 is very similar; the MBS method can be successfully used on serotype 3 samples, which are of particular interest due to the reduced effectiveness of PCV13 on serotype 3 IPD.^24–26^

We demonstrate that good separation can be achieved with only one unique antiserum, meaning that serotypes with cross-reactivity to one antiserum can still be separated using this method. As expected, we demonstrate that the efficiency of enrichment achieved by each of the two antisera pools is not equal and therefore, depending on the desired serotype one antisera may be preferred over another. Furthermore, enrichment of a serotype can occur even when a serotype is present at only 0.01% of the total sample (1×10^3^ minority serotype with 1×10^7^ majority serotype).

A key limitation of the MBS method in general, is that due to cross-reactivity within serogroups, SSI antisera Pools can only be used to separate *S. pneumoniae* serotypes belonging to different serogroups. However, use of type-specific antisera or a mAb instead of pooled antisera would circumvent this limitation. Another limitation is the total proportion of minority cells that can be recovered. While enrichment from 0.1% up to >10% has been demonstrated, it is worth noting that only a small proportion (∼1%) of the total minority cells present in the initial mixture are successfully extracted. This may be overcome by increasing antibody incubation periods or antibody concentration to increase binding capacity.

Having optimized the MBS method, we evaluated its potential for laboratory applications. The MBS method allows for higher throughput generation of capsule-switch variants, by combining the method with the existing techniques, such that gDNA from multiple donor serotypes can be transformed into a recipient serotype in a single mixed reaction. After initial selection for transformants on selection media, the MBS method can be used to separate out the individual transformants in a serotype-specific manner. Mixed transformations would permit higher throughput generation of capsule-swapped variants, the potential to determine comparative efficiency and a significant reduction in BAP usage and labor intensity. However, in the absence of MBS, whilst isolation of different serotypes is comparable to that observed in individual transformations, the benefits are offset by the lengthy and time-consuming process of serotype screening each isolate by latex agglutination. Therefore, to harness the true benefit of mixed transformations a simple and easy technique to select for different serotypes is required. The MBS method was used to isolate multiple serotypes, from a mixed sample of four serotypes. The MBS method outperformed the individual transformations and the mixed transformation (without MBS), by successfully isolating an additional serotype (11B) which was not isolated using the other methods. This suggests that the MBS method may be particularly useful to enrich for serotypes which transform with low efficiency. The MBS technique was not 100% specific, and a small amount of cross-reactivity was observed, however, since each sample is enriched for the desired serotype, and the serotype of each colony is confirmed by latex agglutination, these contaminants are of little concern are for this particular application.

We also show that the MBS method can be modified to successfully enrich for pneumococci of a known serotype from saliva samples. Enrichment is possible even in saliva samples where pneumococci is present at very low concentrations (5×10^1^ CFU/mL), for which isolation of pneumococci using standard methods is typically very challenging. This permits easy identification and isolation of pneumococci present in saliva at concentrations too low to detect using standard dilution and plating methods. The use of SSI antisera alone on a polymicrobial sample such as saliva was problematic due to antisera reactivity with non-pneumococcal bacteria present in saliva. In general, we found that the SSI antisera outperformed mAbs in terms of total number of pneumococcal colonies isolated, we hypothesize that this is due to the increased avidity of antisera (presence of IgA, IgM) which agglutinates pneumococci increasing the overall yield during MBS. Therefore, to take advantage of the increased avidity of antisera and simultaneously the high specificity of mAbs, we combined both in the primary incubation step, but only targeted the mAb in the secondary antibody step. While the elution from saliva samples was not 100% pure pneumococci, contaminating non-pneumococcal bacteria was reduced, and identification and selection of single pneumococcal colonies was improved when compared to the standard dilution and plating method. The enrichment observed varies depending on concentration of pneumococci present in the sample, but also on the saliva composition itself. The composition of bacterial community in saliva varies between different age groups^27^ and so the success of the MBS method will likely vary accordingly. Since the MBS method can work on saliva containing very low concentrations of pneumococci it may be particularly useful for the isolation of minority serotypes in samples obtained from multiply colonized individuals. Previous research shows that 52% of Dutch primary school children tested positive for multiple pneumococcal serotypes,^19^ however, conventional serotyping methods often result in an underestimation of multiply colonized individuals.^28^ Detection of multiple serotypes is possible using serologic, biochemical (Mass Spectroscopy and nuclear Magnetic Resonance), and genotypic (sequencing, qPCR and microarrays) methods, however, until now, attempting to isolate minority serotypes by conventional methods (single colony selection) has been laborious and time consuming.^21^

## CONCLUSION

The MBS method allows for the successful enrichment of a minority serotype from a dual sample containing two *S. pneumoniae* serotypes belonging to different serogroups. Using this method, an initial sample containing 0.01-0.1% of a desired serotype, can be enriched to up to 90% in the final sample. Enrichment to between 10 and 90% was demonstrated for six minority serotypes, and half of the commercially available antisera pools (Pools B, E, H, P, Q, R and S) were tested. We demonstrate two different applications for this technique: separating capsule-switch variants from mixed transformation experiments and enriching for pneumococci of a known serotype from saliva. The MBS technique can be used successfully to enrich for serotypes which are present at very low-levels in both mixed cultures and more complex polymicrobial sample types (such as saliva), making it a versatile and important technique for a multitude of applications.

## AUTHOR CONTRIBUTIONS

Conceptualization, DMW and ALW; Methodology, AY, DMW and ALW; Investigation, AY, EH, SM, MH, DYC, HE, JR; Analysis, AY and DW; Writing – initial draft, AY; Writing – Review & Editing, DMW, ALW and DYC; Supervision, ALW, DMW, JR.

## AUTHOR DECLARATION

DMW has received consulting fees from Pfizer, Merck, GSK, Affinivax, and Matrivax and is PI on research grants from Pfizer and Merck to Yale. ALW has received consulting fees from Pfizer and is PI on research grants from Pfizer to Yale. The other authors declare no conflict of interest.

## Supplemental Information

**Table S1.** Serotypes and corresponding antisera pools used for MBS (rabbit antiserum; SSI Diagnostica, Hillerød, Denmark) and for serotyping (ImmuLex™ Pneumotest; SSI Diagnostica).

| Serotype | Pooled Antisera for Neufeld | ImmuLex™ Pneumococcus Antisera |
| --- | --- | --- |
| 12F | Pool E #16733<br>Pool R #16741 | Pool E #52394<br>Pool R #52401 |
| 23F | Pool H #16736<br>Pool Q #16740 | Pool H #52397<br>Pool Q #52400 |
| 3 | Pool B #16728<br>Pool R #16741 | Pool B #52391<br>Pool R #52401 |
| 14 | Pool P #16739<br>Pool H #16736 | Pool P #52399<br>Pool H #52397 |
| 19A | Pool B #16728<br>Pool P #16739 | Pool B #52391<br>Pool P #52399 |
| 15A | Pool H #16736<br>Pool S #16742 | Pool H #52397<br>Pool S #52402 |
| 2 (D39) | Pool A #16725<br>Pool T #16743 | Pool A #52390<br>Pool T #52403 |
| 35B | Pool G #16735 | Pool G #52396 |
| 11B | Pool D #16731<br>Pool T #16743 | Pool D #52393<br>Pool T #52403 |

**Table S2.**
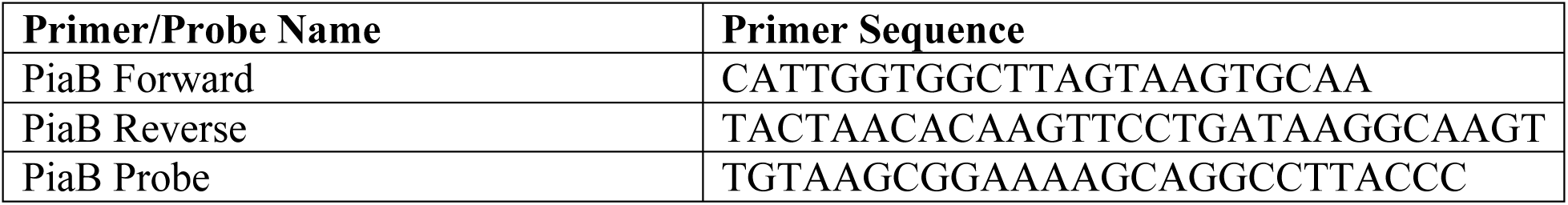
PiaB qPCR primers and probes.

**Table S3.**
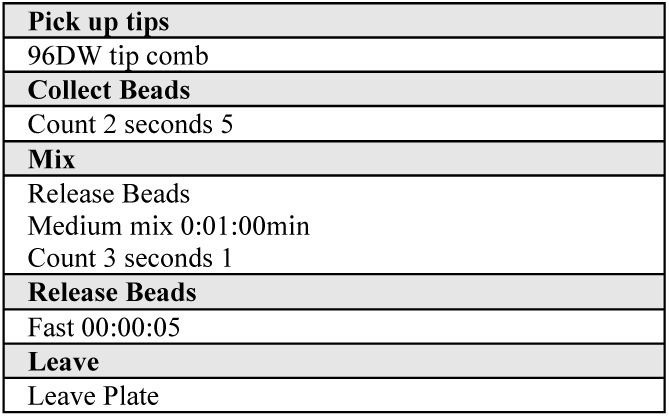
KingFisher Flex MBS Protocol

**Table S4.**
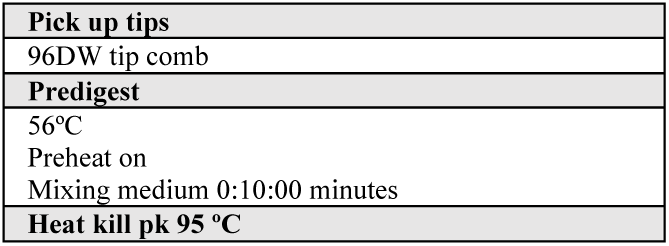
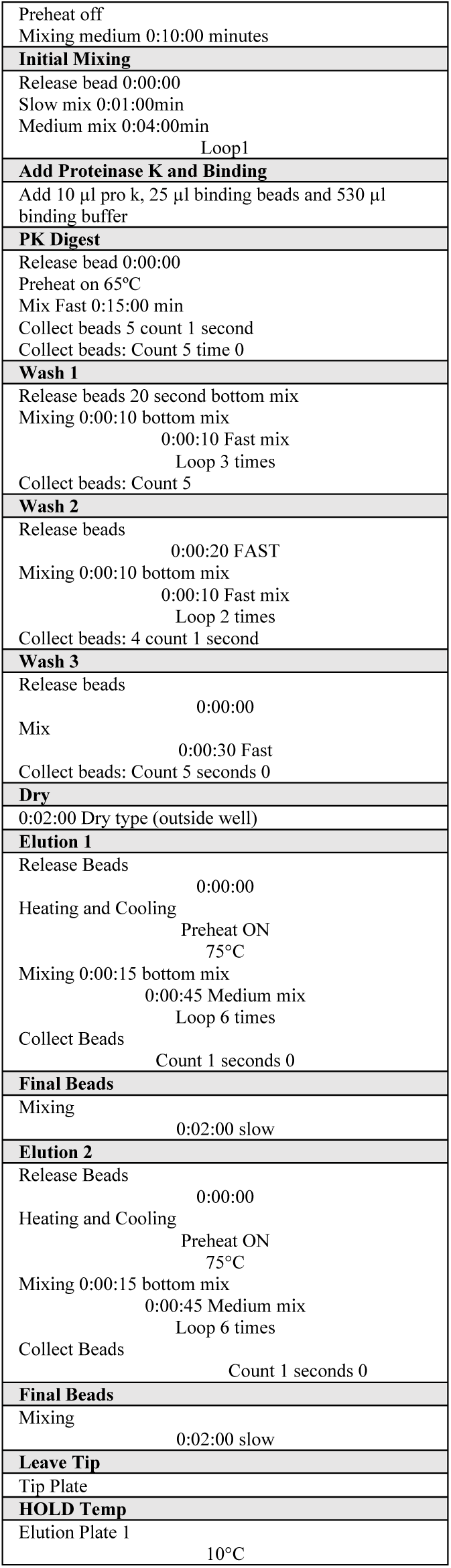
KingFisher Flex DNA extraction protocol

**Figure S1.**
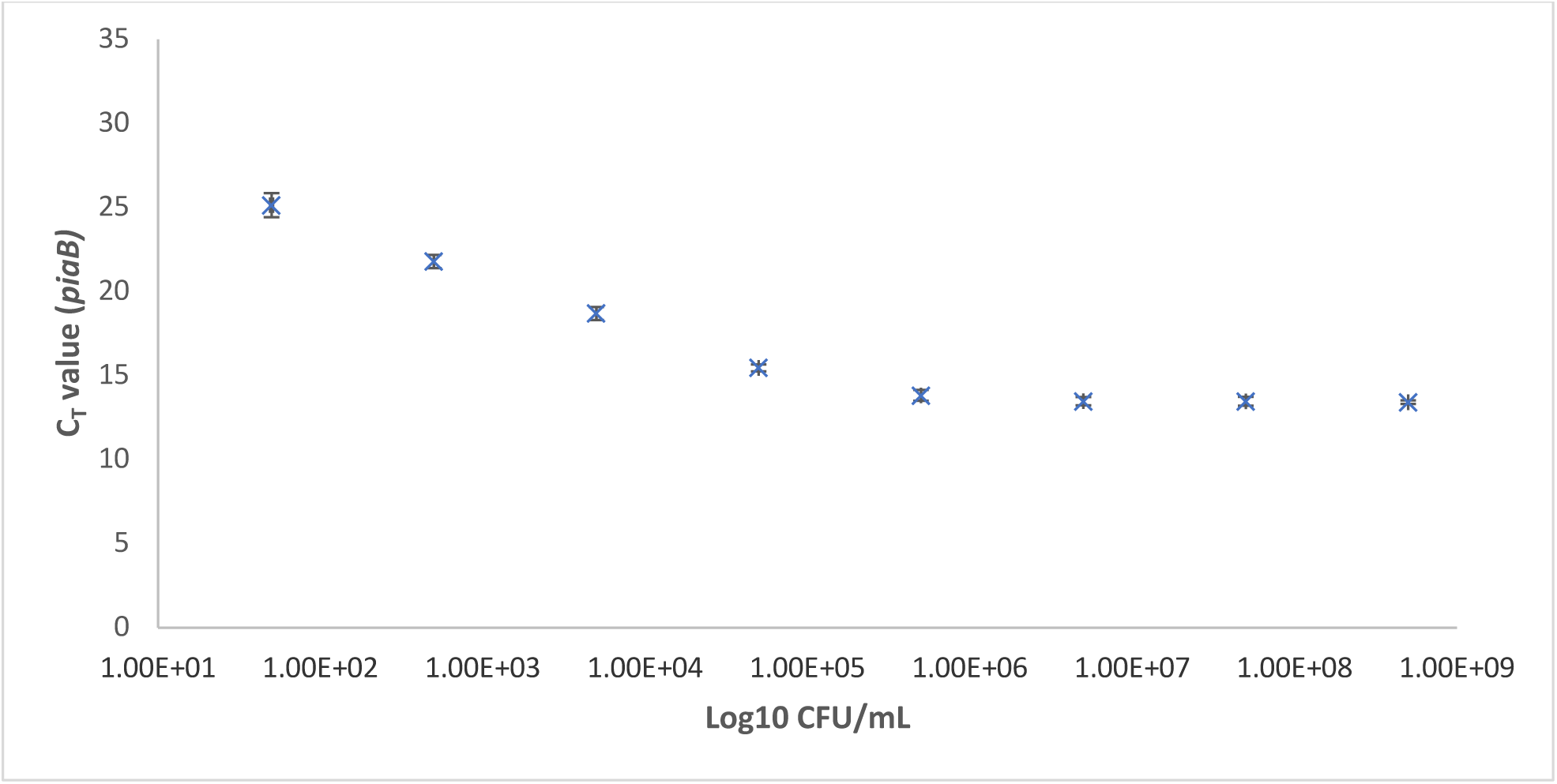
Detection of pneumococcus gene piaB (C_T_ value) when culture-enriched saliva samples were tested with qPCR, and the corresponding CFU/mL of S. pneumoniae 19A that was spiked into each raw saliva sample. Raw saliva was confirmed to be pneumococcus-negative (C_T_ >40) by qPCR towards piaB. Data shown as mean and standard deviation of biological triplicate data.

## Supplemental File - Raw Data

